# Transposon-Display of AI-designed binders enables manipulation of the proteome in human cells

**DOI:** 10.1101/2025.05.11.653342

**Authors:** Alexander Hoch, Caroline I. Fandrey, Brian Coventry, Sebastian Virreira Winter, Philipp E. Geyer, Tim N. Kempchen, Thomas S. Ebert, Michael Hölzel, David Baker, Jonathan L. Schmid-Burgk

## Abstract

Transposon-Display is a highly scalable screening method that links proteins to their encoding DNA during expression in *E. coli* via a mutant transposase. Leveraging this system, we identified AI-designed binders targeting four intracellular human proteins, saturation-mutagenized top candidates, and charge-balanced variants to improve compatibility with the intracellular environment while preserving target binding. The resulting neutral binders can be fused to EGFP allowing antibody-free intracellular staining. Expressing neutral binders in human cells as fusions with functional domains can drive small molecule-controlled protein aggregation and trigger proteasomal degradation of endogenous target proteins in living cells. Manipulating the proteome of living cells using libraries of AI-designed binders may provide a new avenue to screen for disease-relevant protein functions and large-scale functional data may help to refine protein design algorithms.

## INTRODUCTION

An outstanding challenge in biology is the manipulation of intracellular proteins to study their function. While single-chain antibodies such as scFv fragments or camelid nanobodies can be expressed in human cells as intrabodies to fluorescently label or degrade specific proteins ^1^, developing these reagents typically requires elaborate methods like camelid immunization or high-throughput library screening ^2 3^. Computationally designed binder proteins provide a promising alternative due to their small size, stability, and encodability in library formats ^4 5^.

Structure-generating diffusion models such as RoseTTAFold diffusion (RFdiffusion) enable de novo design of protein scaffolds with tailored binding interfaces ^6^, and deep learning sequence-optimization tools like ProteinMPNN can rapidly generate sequences that reliably fold and bind as intended ^7^. Despite these innovations, computational designs often require refinement to achieve the high affinities and robust function needed in living cells. In the natural immune system, antibodies achieve high affinity through iterative mutation and selection (affinity maturation); analogously, initial AI-designed binders can be further optimized via directed evolution in the laboratory ^4^.

AI-designed proteins open the door to a wide range of new applications, for example, the precise interference with signaling pathways by selectively blocking or degrading endogenous proteins in living cells. Unlike RNAi or CRISPR-based approaches, which act at the transcript or genome level, AI-binders may enable time-controlled direct, reversible, and isoform-specific manipulation of protein function - ideal for dissecting dynamic cellular processes. Furthermore, AI-binders could be fused to effector domains to create modular tools for proximity labeling, targeted ubiquitination, or localization control. Their small genetic footprint makes them amenable to delivery via viral vectors, facilitating applications in hard-to-transfect cells and in vivo models. Ultimately, the combination of AI-guided design and rapid experimental optimization may enable the creation of a new class of programmable protein tools, capable of targeting virtually any intracellular protein with high specificity and minimal off-target effects.

To facilitate this, we introduce a novel display technology that is built on a transposase protein. We demonstrate rapid enrichment and optimization of AI-generated binder designs targeting immune-relevant proteins and their use analogous to antibodies in staining and pull-down applications. Moreover, we modulate the proteome of live cells using AI-binder fusion proteins.

## RESULTS

### High-throughput binder screening using optimized transposase mutants

DDE transposases recognize DNA sequences referred to as inverted terminal repeats (ITRs) to excise and bind their own encoding DNA. In their natural context, ITRs are marking the boundaries of mobile genetic elements, which can thus be inserted into a new genetic locus. Based on a mutant *PiggyBac* transposase ^8^ that can excise and bind but not reintegrate ITR-flanked DNA, we devised a binder screening method that couples a library of proteins to their own encoding DNA in *E. coli* (Fig. 1a). Upon lysing the bacteria, protein-DNA complexes can be enriched by affinity pulldown, and bound DNA can be sequenced by NGS to identify high-affinity binders. As an initial test of this approach, we fused peptides and nanobodies with known binding partners to the transposase in different configurations and quantified their target-specific enrichment by sequencing (Fig. 1b). While some configurations displayed an up to 1,000-fold enrichment, non-binding sequences were not fully eliminated even by extended washing. To improve the system’s enrichment rate of correct binders from very large libraries, we profiled single amino acid mutations throughout a large fraction of the transposase protein, fused to a FLAG peptide that can be enriched using antibody pulldown. Additionally, we introduced three silent mutations around each randomized codon to reliably distinguish true mutations from sequencing errors (Fig. 1c). As expected, mutations in most regions of the transposase that generate a stop codon diminished sequence enrichment, whereas stop codons in the C-terminal domain starting at residue 540 of the transposase consistently improved enrichment by as much as 20-fold at some positions (Fig. 1d). Furthermore, several single amino acid mutations increased the enrichment ratio by >3-fold. To validate the improved enrichment characteristics, we randomized four out of eight consecutive codons of a FLAG peptide fused to wild type or mutant transposase and screened for binders to a commonly used anti-FLAG antibody, confirming an >8-fold improvement of the C-terminally truncated *PiggyBac* transposase (Fig. 1e). Peptides containing a DYK motif were enriched by several thousand-fold from a pool of >100,000 sequences in a single round of selection (Fig. 1f). We name this pooled binder screening method Transposon-Display.

**Figure 1:**
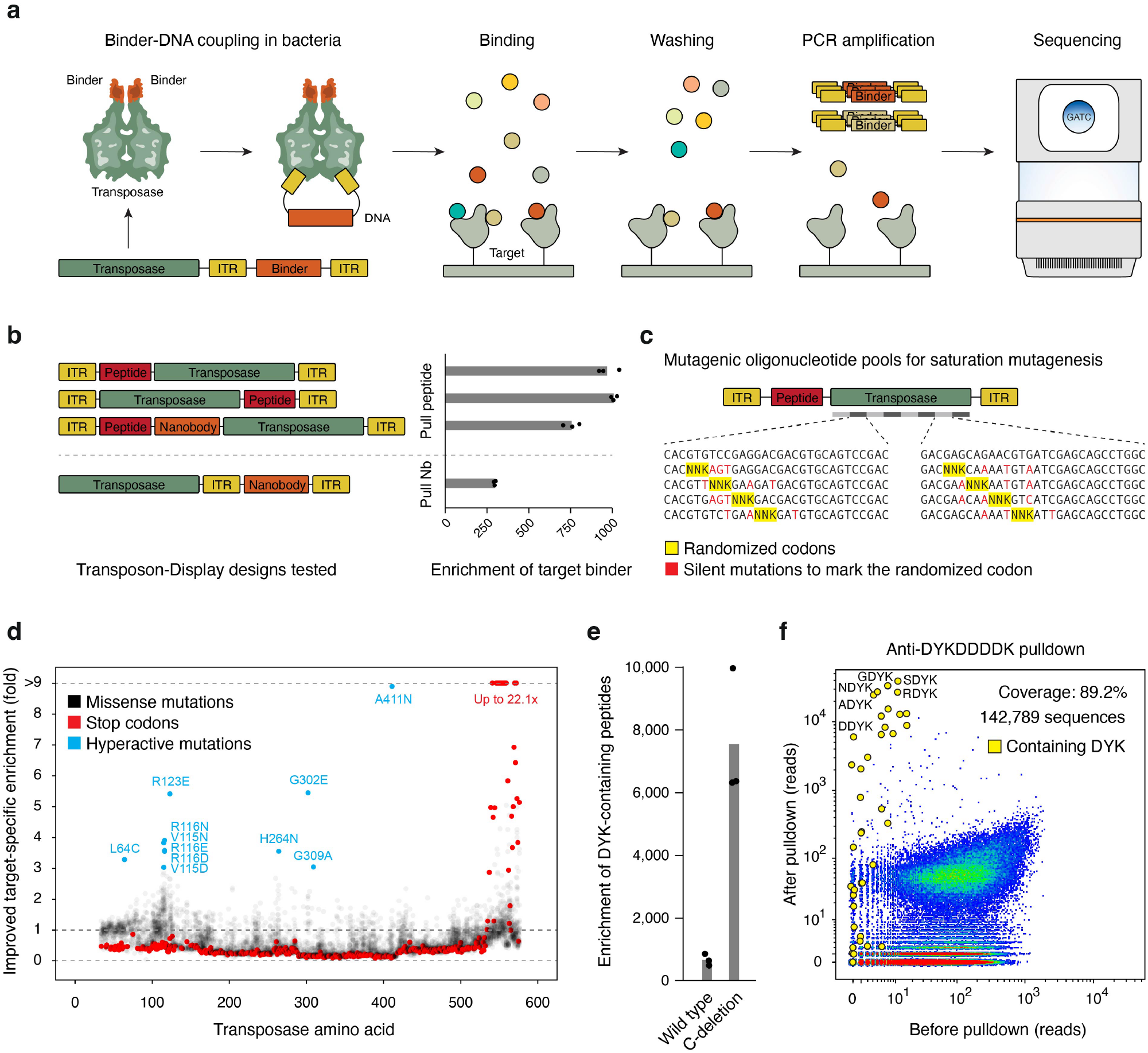
Optimization of Transposon-mediated screening of biomolecules. **(a)** Schematic of Transposon-Display. A transposase fused to a biomolecule links the biomolecule to its encoding DNA. Upon affinity-based enrichment of biomolecules from a pool, DNA sequencing is used to identify strong binders. **(b)** Sequence configurations tested for Transposon-Display, using FLAG peptide or an EGFP-binding nanobody as exemplary biomolecules fused to *PiggyBac* transposase. **(c)** Schematic of saturation mutagenesis of *PiggyBac* transposase to identify optimized mutants. The transposase coding sequence was tiled by oligonucleotide pools of around 100 nucleotides in length, covering each codon with an individual NNK-randomized oligonucleotide. Unique silent mutations are introduced into each oligo to enable the reliable identification of mutations from deep sequencing data. **(d)** Results of saturation mutagenesis of *PiggyBac* transposase. Mutants enriched after pulldown are highlighted. While stop codons were deleterious throughout most of the transposase gene, truncations of the C-terminus around residue 550 enabled >20-fold higher enrichment by Transposon-Display. **(e)** Validation of the enhanced transposase variant in the context of screening a four-amino acid peptide library by antibody pulldown. DYK-containing peptides are enriched at increased rates by the truncated transposase. **(f)** Pooled Transposon-Display screening of a randomized peptide library binding to an anti-FLAG antibody. Most enriched sequences display strong similarity to the known binding motif of the antibody.

### Transposon-mediated identification and optimization of AI-designed affinity reagents targeting human intracellular proteins

To enable rapid identification of AI-designed affinity reagents and optimize their affinity, we modified the Transposon-Display vector to enable Golden Gate cloning of thousands of AI-binders from oligonucleotide pools as C-terminal transposase fusions. Codons were randomized to enable binder identification by short sequencing reads while excluding rare codons and restriction sites interfering with Golden Gate library cloning. For pooled binder screening, human target proteins were transiently over-expressed as EGFP-3xFLAG-fusions in human HEK 293T cells, and crude lysate was bound to anti-FLAG magnetic beads (Fig. 2a). AI-binder Transposon-Display pools were produced in *E. coli* and bound to the same beads. Binder distributions were determined by read counting before and after bead pulldown, which confirmed high coverage across four human targets (Fig. 2b). Between 2 and 41% of 500 designs for each target protein were identified as true binders (Fig. 2c). For affinity maturation of AI-binders, we performed saturation mutagenesis using oligonucleotide pools covering up to 33 amino acid regions with NNK degenerate codons at every possible position, flanked by three silent mutations to enable reliable identification of the mutated position from sequencing data (Fig. 2d). Exemplary results from mutagenizing an AI-binder for human PYCARD yielded a 100% coverage of possible mutations, half of which abolished the binding properties of the molecule (Fig. 2e) while multiple mutations at hotspot position lysine 44 as well as several point mutations increased binder enrichment by Transposon-Display by up to 14.7-fold (Fig. 2e). Of note, numerous mutations were identified that do not affect binding yet neutralize or invert the negative charge of amino acid residues. In contrast to high-affinity single-chain antibodies, most AI-binder designs were predicted to be negatively charged at pH 7.4. To adjust their predicted charge to −0.5 – +0.5 at pH 7.4, we screened random combinations of neutralizing mutations in combination with the high-affinity mutation K44S using Transposon-Display and identified multiple charge-neutral high-affinity binders (Fig. 2f). Crude lysates of bacteria expressing neutral binders fused to EGFP stained endogenous PYCARD (ASC) inflammasome specks in the cytosol of PFA-fixed and permeabilized human primary macrophages dependent on inflammasome activation by PrgI+PA stimulation, while a non-optimized binder failed to produce stimulation-specific staining signal (Fig. 2g). To assess specificity, we immobilized an AI-binder fused to EGFP on anti-EGFP agarose beads and pulled binding proteins from the lysate of phorbol 12-myristate-13-acetate (PMA)-differentiated LPS-stimulated human THP1-derived macrophages. Proteomics by data-independent acquisition mass spectrometry confirmed the expected target protein PYCARD as the top-enriched protein as compared to EGFP-only control beads (Fig. 2h).

**Figure 2:**
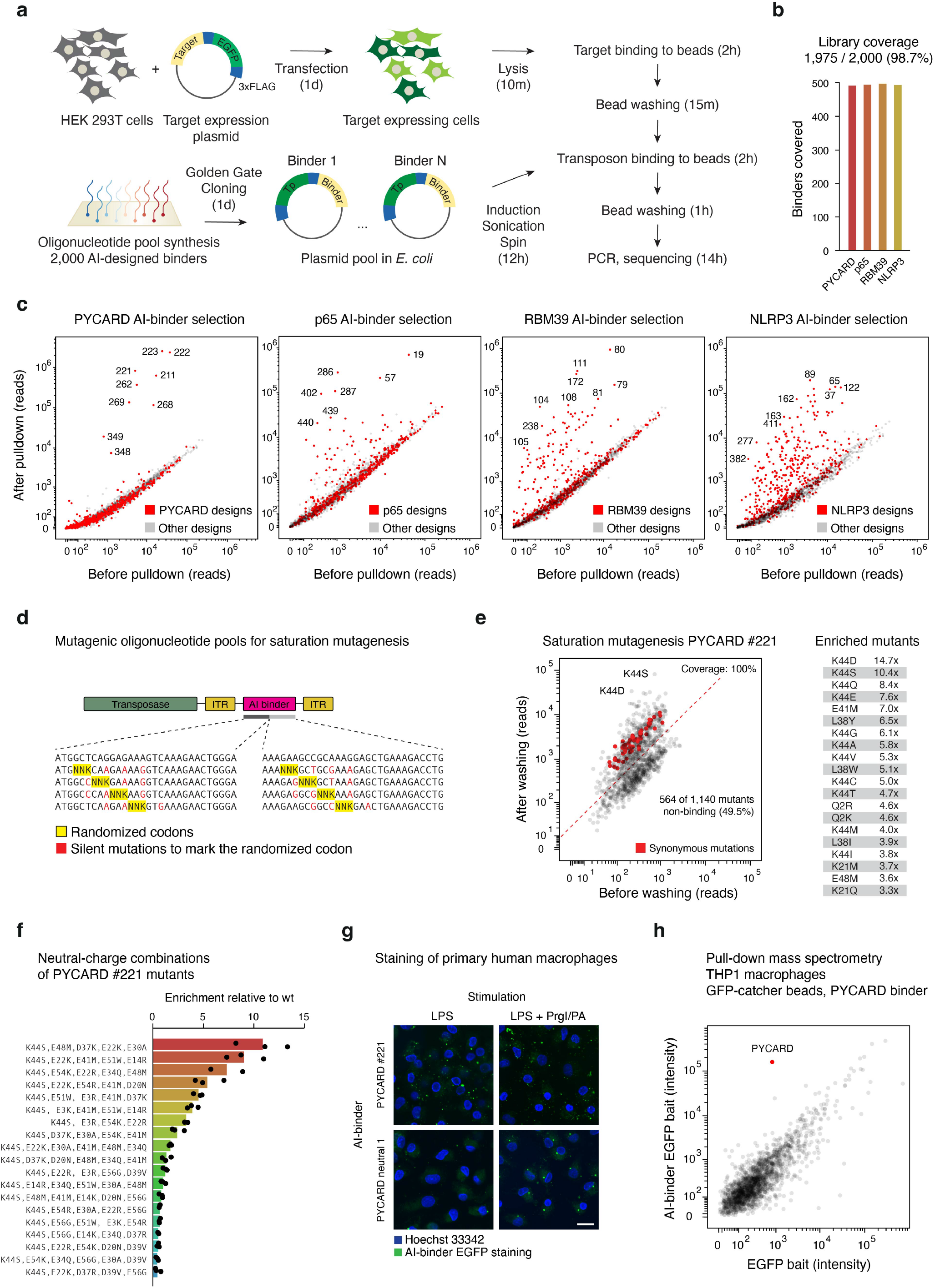
Selection and optimization of AI-designed binders targeting human proteins. **(a)** Schematic of Transposon-Display of AI-designed binders for intracellular human targets. Target proteins are expressed as EGFP-3xFLAG-fusions by transient transfection of HEK 293T cells. AI-binder libraries are generated using RFdiffusion, reverse translated using diversified codons, synthesized as oligonucleotide pools and inserted into a Transposon-Display plasmid using Golden Gate cloning. Screening can be completed in three days. **(b)** Library coverage after pooled cloning of 2,000 AI-binders targeting four intracellular human proteins, as quantified by NextSeq 2000 sequencing. **(c)** Results of Transposon-Display screening of 500 on-target AI-binders targeting four intracellular human proteins (red dots) in a pool with 1,500 non-targeting AI-binders (grey dots). **(d)** Saturation mutagenesis of AI-designed binders to identify improved binder mutants. The AI-binder coding sequence was tiled by oligonucleotide pools of up to 100 nucleotides in length, covering each codon with an individual NNK-randomized oligonucleotide. Three silent mutations are introduced into each oligo to enable the reliable identification of mutations from deep sequencing data. **(e)** Results of saturation mutagenesis of an AI-designed binder targeting human PYCARD. Mutants enriched or lost after pulldown are highlighted. Top-enriched mutants are listed with their fold-enrichment over the wild-type binder. **(f)** Transposon-Display results of combinations of mutations selected from (e). Random combinations of mutations were chosen to even out the predicted charge at pH 7.4 to between −0.5 and 0.5. One of the strongest mutations from (e), K44S, was included in all designs. Displayed are relative enrichment scores over the wild-type AI-designed PYCARD binder #221. **(g)** M-CSF differentiated macrophages from the blood of a healthy human donor were stimulated with LPS or LPS/PrgI+PA to activate the NLRC4 inflammasome and PYCARD speck formation. Cells were fixed with PFA, permeabilized with Triton X-100, and stained with unpurified lysate of *E. coli* expressing AI-binder EGFP fusions. After washing the cells three times with PBS and staining nuclear DNA with Hoechst 33342, images were acquired by spinning-disc confocal microscopy. The expected staining pattern upon PrgI/PA-mediated inflammasome activation is a single PYCARD speck per cell localized close to the nucleus. **(h)** AI-binder EGFP fusions were expressed in *E. coli*. After sonication and centrifugation, the binders were captured on GFP-Catcher beads. Lysates of PMA-differentiated THP1 macrophages were incubated with binder-loaded beads. Bound proteins were digested with trypsin and analyzed via mass spectrometry.

### Drug-induced protein aggregation and proteasomal degradation in human cells

To test whether AI-binders can be used to manipulate the proteome in living cells, we first turned to the BTB domain of Bcl6, which aggregates in response to a cell-permeable small molecule ^9^. This domain was fused to validated AI-binders and mCherry and expressed in human cells which co-express the EGFP-tagged target protein (Fig. 3a). Upon addition of BI-3802, mCherry formed an aggregate in positively transduced cells, and EGFP-tagged proteins co-aggregated with a target-matching AI-binder (Fig. 3b). We conclude from this experiment that the two neutral-charge AI-binders tested successfully bind their target protein in the context of the cellular cytosol when expressed via single-copy lentiviral integration. To test whether AI-binders also enable targeted degradation of proteins, we fused them to a mutated FKBP domain for which small molecule PROTACs have been developed ^10 11^ (Fig. 3c). Expressing AI-binder fusions through lentiviral delivery in human cells which co-express EGFP-tagged target protein and an mCherry marker for normalization, we found most cells to transition from an EGFP mCherry double-positive to an EGFP-negative gate in a binder-dependent manner upon stimulation with the PROTAC dTAGV-1 (Fig. 3d). Finally, to test endogenous protein degradation we transduced human THP1 monocytes with lentivirus to express AI-degrader constructs. After selecting for positively transduced cells using Puromycin, cells were differentiated to macrophages, LPS-primed, and monitored for PYCARD protein expression upon PROTAC stimulation by Western blot. While neither the PROTAC nor the AI-binders alone affected endogenous PYCARD expression levels, both neutral-charge AI-degraders abrogated protein expression in a PROTAC-controlled manner (Fig. 3e).

**Figure 3:**
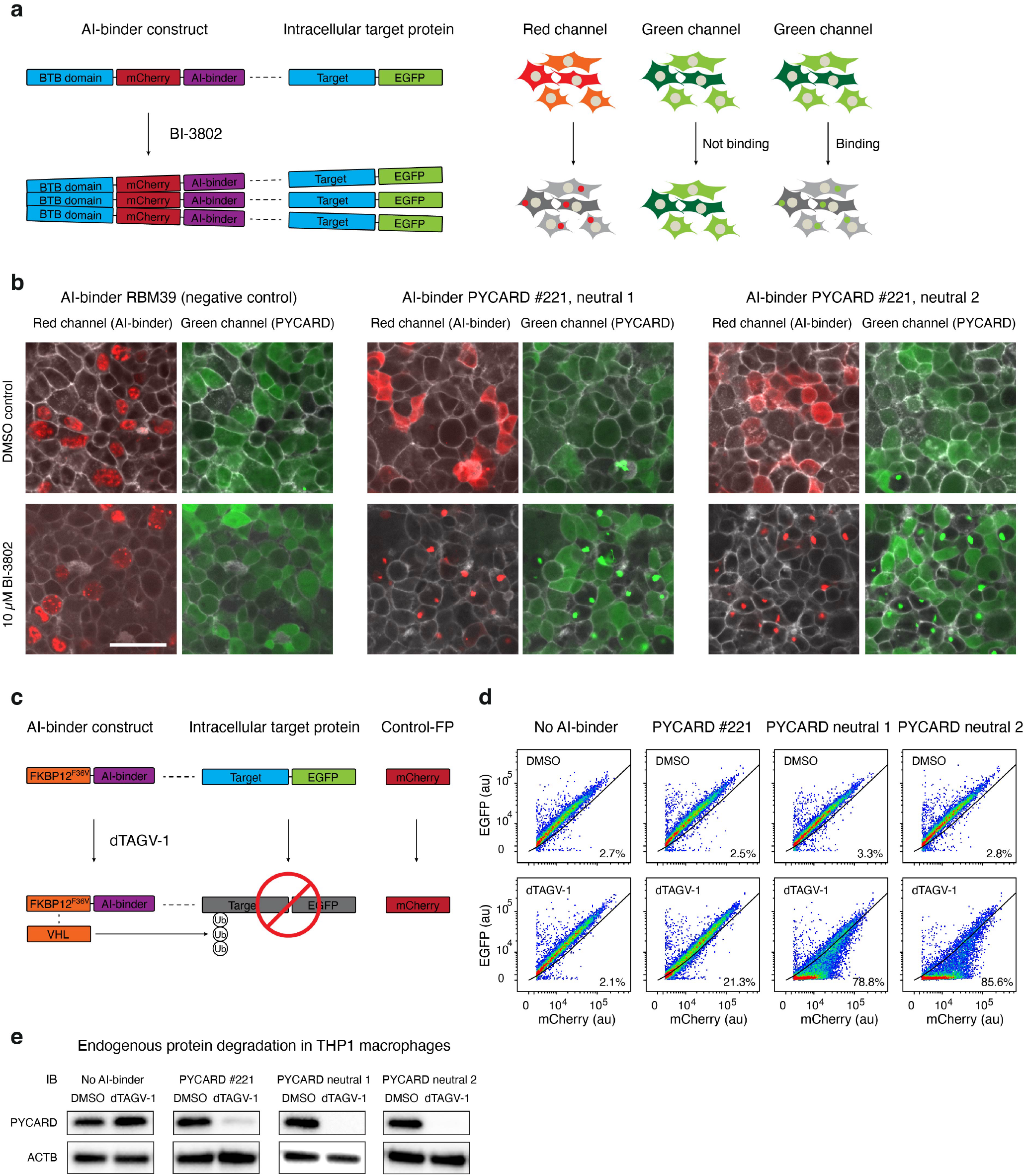
Application of optimized AI-binders to manipulate the proteome of human cells. **(a)** Principle of fusing AI-binders to a BTB domain derived from human Bcl6, which aggregates in live cells upon addition of the small molecule BI-3802. By fusing fluorescent proteins to both the AI-binder and the target protein, target binding of the AI-binder can be monitored using live fluorescence microscopy. **(b)** HEK 293T cells were transduced with a lentivirus to express EGFP-fused PYCARD. Cells were sorted for EGFP expression. Two improved variants of human PYCARD binders as well as an RBM39-binding control were expressed as fusions to a BTB-domain and mCherry through a second lentiviral transduction. Images show drug-induced aggregation of AI-binders, and co-aggregation of EGFP upon target binding. **(c)** Principle of fusing AI-binders to a mutant FKBP12 domain, which can recruit an E3 ubiquitin ligase using the small-molecule PROTAC dTAG^V^-1 and induce proteasomal degradation of the bound target protein. By fusing a fluorescent protein to the target protein and co-expressing a second fluorescent protein for normalization, targeted protein degradation can be quantified using flow cytometry. **(d)** HEK 293T cells were transduced with a lentivirus expressing EGFP-fused PYCARD and mCherry separated by a 2A-peptide. Either an AI-binder targeting PYCARD or neutral variants of the same binder were expressed as fusions to an FKBP12^F36V^-domain through a second lentiviral transduction, for which cells were selected using Puromycin. Plots show flow cytometry-based quantification of targeted protein degradation across single cells upon incubation with 10 µM dTAG^V^-1, gated for mCherry expression. **(e)** THP1 monocytes were transduced with a lentivirus expressing AI-binders fused to an FKBP12^F36V^-domain. Puromycin-selected cells were differentiated, LPS-primed, and treated with 1 µM dTAG^V^-1 or DMSO for 24 hours. Western blots of cell lysates were stained with anti-PYCARD and anti-ACTB antibodies and developed using HRP detection.

## DISCUSSION

Over the past four decades, display technologies have fundamentally shaped the field of molecular biology, beginning with the invention of phage display by George P. Smith in 1985 ^12^. Since then, a diverse array of methodologies like ribosome display ^13^, mRNA display ^14^, and yeast surface display ^15^ have been developed, each offering unique advantages for the selection of high-affinity binders. Transposon-Display introduces an approach that utilizes a highly compact genotype– phenotype linkage among current display systems by coupling a *PiggyBac* transposase to a binder protein and the same binder’s coding DNA sequence. With three days of hands-on time, low resource and safety requirements, it can be performed in any molecular biology lab using only basic equipment. A truncated transposon variant routinely achieved enrichment rates of >5,000-fold in a single round of screening, which is comparable to the high enrichment rates of multiple phage display rounds ^16^.

Applying the Transposon-Display workflow, we enriched and optimized AI-designed protein binders targeting immunologically relevant intracellular proteins. This workflow could be scaled up for creating a public resource of validated affinity reagents as an alternative to antibodies. Transposon-barcoded AI-binders could furthermore enable high-multiplex DNA-based tissue staining using CODEX-like protocols ^17^, in which expensive antibodies are replaced by AI-binders produced and barcoded in a pool. DNA identifiers bound to multiple AI-binders targeting separate interfaces of the same target protein may furthermore increase the specificity in diagnostics applications through proximity ligation or proximity elongation analogous to methods like NULISA^18^. To that end, the transposon-bound dsDNA could be cleaved or partially linearized, enabling free DNA strands of adjacent binders to anneal for subsequent ligation or polymerase-mediated elongation and identification via sequencing. Transposon-Display AI-binder pools could also be used as a multiplexed protein readout in single cell sequencing applications, translating the presence of proteins into DNA barcodes analogous to PHAGE-ATAC^19^.

Mass spectrometry and intracellular staining results demonstrate that AI-binders can bind their target protein selectively in a complex cellular context. By expressing AI-binders coupled with functional protein domains, we were able to aggregate and degrade specific proteins in living human cells. Compared to previous studies, which directly added degrons to target proteins ^20^, fused nanobodies to a degron domain ^21^, or used antibodies with the Fc receptor TRIM21 ^22^ to degrade target proteins, AI-degraders may provide a much simpler strategy for targeted protein degradation. Given the small size of AI-binders, this approach of manipulating the proteome might enable pooled functional screening using lentiviral delivery, analogous to expressing libraries of gene-targeting sgRNAs in CRISPR screening applications.

Functional screening at the level of proteins provides multiple hypothetical advantages, which are 1) a higher kinetic resolution due to circumventing transcription and translation, 2) the ability to transiently inhibit essential proteins, 3) the inhibition of individual protein surfaces, isoforms, or protein-ligand interactions, 4) more specific ways to interfere with protein function, like re-localizing, aggregating, or gluing proteins together, and 5) not requiring the delivery of large CRISPR nucleases, which limits screening applications in hard-to-transduce or short-lived cell types.

Transposon-Display provides a simple approach to identify and affinity-mature AI-binders interfering with specific proteins in human cells. At the same time, the refinement of computational design and scoring algorithms for AI-binders may eventually make physical binder screening obsolete. While current state-of-the-art interface scoring models like Chai-1 ^23^ partially predict binder strength (Fig. 4a-b), collecting large-scale Transposon-Display datasets of successful and failed AI-binder designs could provide a key resource for training more accurate binder design and affinity prediction algorithms in the future.

**Figure 4:**
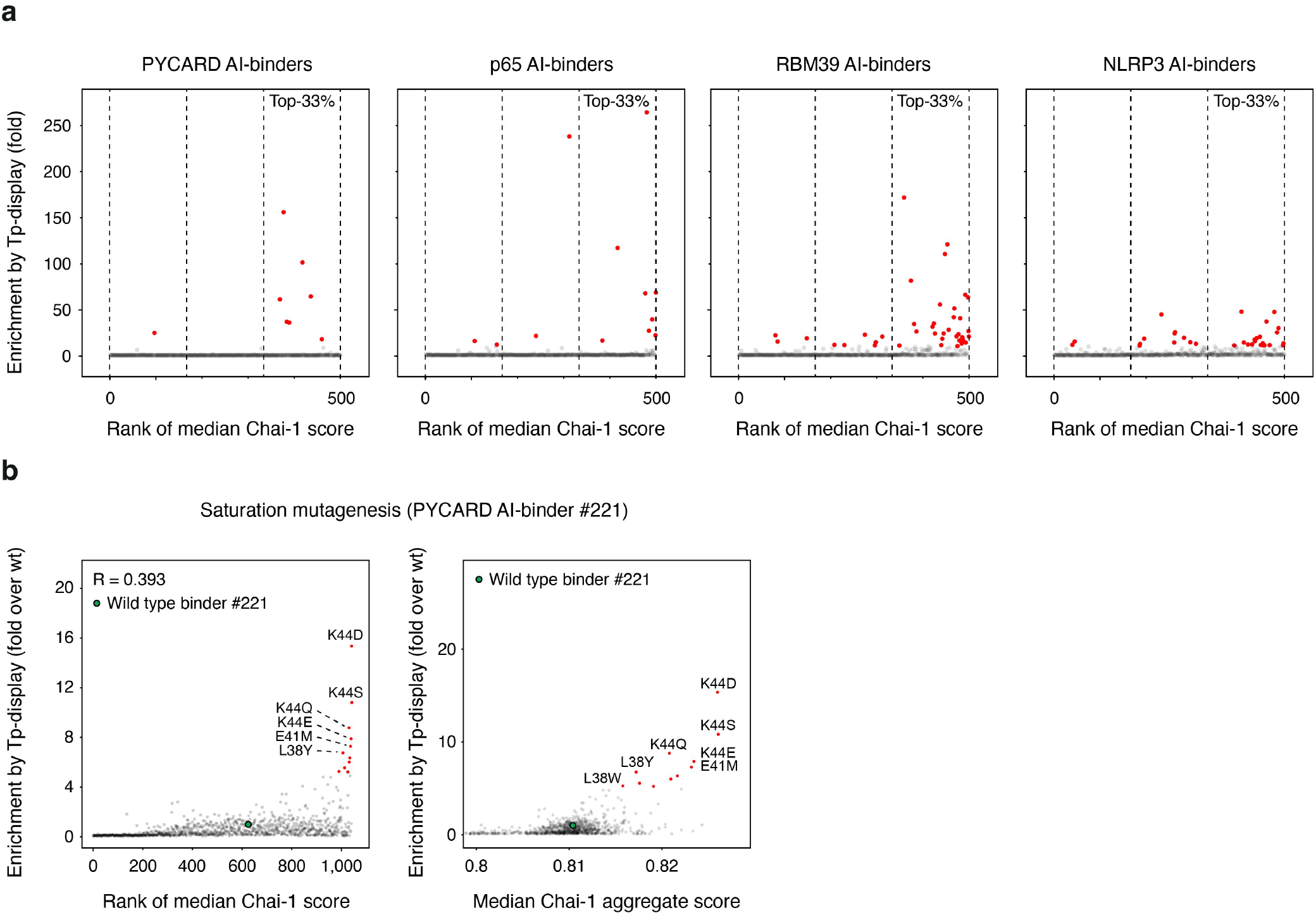
Enrichment of best-performing binders by Chai-1 aggregate scores. **(a)** For each target, 500 AI-designed binders were co-folded with their target domain using Chai-1 in five replicates. The AI-designs were ranked by the median of the five aggregate scores. Plots depict Transposon-Display read enrichment rates as a proxy for binding strength in relation to their predicted rank. **(b)** Single amino acid mutations of an AI-designed binder for human PYCARD were co-folded with their target domain using Chai-1 in five replicates. The mutants were ranked by the median of the five aggregate scores. Plots depict Transposon-Display enrichment normalized to the wild type binder in relation to their predicted rank (left panel) or to their median aggregate score (right panel).

## Acknowledgements

We thank Florian Schmidt for providing the ASC-GFP expressing HEK 293T cell line and Florian Schmidt and Eicke Latz for primary macrophages. We thank Marta Lovotti, Peter Konopka, David Feldman, Marius Jentzsch, Katja Blumenstock, Maia Cristodaro, Gregor Hagelueken, Stephan Menzel, Radosław Nowak, Jürgen Bajorath, Tobias Bald, Eicke Latz, and Rayk Behrendt for helpful discussions. We thank the FACS Core Facility of the Bonn Technology Campus (BTC). J.L.S.-B. and M.H. are members of the Cluster ImmunoSensation supported by the Deutsche Forschungsgemeinschaft under Germany’s Excellence Strategy EXC2151 - project ID 390873048. J.L.S.-B. was supported by the Hans und Ria Messer Stiftung. M.H. was supported by the Deutsche Krebshilfe (German Cancer Aid) project grant 70114292 (excellence program for established scientists) and within SFB 1399 by the Deutsche Forschungsgemeinschaft (DFG) – Project ID 413326622. M.H. is a member of the CANTAR project, which receives funding from the Netzwerke 2021 program, an initiative of the Ministry of Culture and Science of the State of North Rhine-Westphalia.

## Author contributions

J.L.S.-B. conceived of the study. A.H., C.I.F., B.C., T.S.E, D.B., and J.L.S.-B. designed the experiments. A.H., C.I.F., S.V.W and P.E.G. collected the data. A.H., C.I.F., T.N.K., M.H., and J.L.S.-B. analyzed the data. A.H., C.I.F, T.S.E and J.L.S.-B. wrote the manuscript with input from all coauthors.

## MATERIALS AND METHODS

### Four random amino acid library cloning

The 4X-FLAG oligo pool was PCR-amplified with 4X-FLAG_fwd/rev primers. Subsequently, the purified PCR product was cloned into the target vector using Golden Gate assembly. The assembled reaction was purified via a Zymo DNA Clean & Concetrator-25 column. The purified library was electroporated into Endura electrocompetent cells (Lucigen), using 50-100 ng/µl DNA and 50 µl cells in 0.1 cm cuvettes (BioRad) at 1800 V, 10 µF and 600 Ω. Cells were directly recovered in 975 µl pre-warmed recovery medium and incubated for 1 h at 37°C and 300 rpm shaking. Then, each culture was transferred to 500 ml LB medium containing 100 µg/ml ampicillin and grown overnight at 37°C 220 rpm shaking. The DNA was purified with a QIAprep Spin Miniprep Kit.

### Transposon-Display

Cloned libraries were re-electroporated using 50-100 ng of cloned and prepped DNA, as previously described. After bacterial cultures were grown overnight, they were diluted 1:100 and grown to an OD600 of 0.4. To induce transposon expression, Anhydrotetracycline (Biomol) was added to a final concentration of 1.33 µM and the bacteria was shaken overnight at 25°C. Bacterial cultures were pelleted in a precooled centrifuge (Eppendorf 5920R) for 45 minutes and resuspended in 10 ml of cold PBS (Mg+, Ca+) with protease inhibitor. To release the transposon complexes, the samples were sonicated on ice with 10 cycles alternating between 10 sec of amplitude 30 pulses and 20 sec pauses (Qsonica Q700). To pellet the bacterial debris, the samples were centrifuged for 30 minutes in a precooled centrifuge and the supernatant was transferred to a new reaction tube, while storing 50 µl of for sequencing.

Furthermore, 40 µl of Anti-FLAG magnetic beads (Millipore M8823) were washed three times in PBS using a magnetic rack (Invitrogen DynaMag-2 Magnet). The beads were added to the lysate and rotated at 4°C for 2 h. Afterwards, the beads were pelleted by centrifuging 1 min and washed three times before storing the beads in 40 µl buffer at −20°C.

### Library preparation and NGS

The magnetic beads and bacterial lysates were directly added to a PCR reaction using ITR_fwd/rev primers with AI binder libraries and 4xFLAG_seq_fwd/ITR_rev with 4xFLAG libraries and NEBNext PCR polymerase. In a second barcoding PCR, a sample specific DNA barcode and sequencing handles were added. Samples were sequenced on an Illumina Miseq and Illumina NextSeq 2000 using a P1 100-cycle cassette.

### Mammalian cell culture

HEK 293T cells were cultivated in Dulbecco’s modified Eagle medium (DMEM) supplemented with 10% (v/v) fetal calf serum (FCS) and 10 μg/ml Ciprofloxacin in a 37°C incubator with 5% CO2. THP1 cells were cultivated in RPMI supplemented with 10% (v/v) fetal calf serum (FCS) and 10 μg/ml Ciprofloxacin in a 37°C incubator with 5% CO2.

### Target protein expression

0.8 × 10^7^ HEK 293T cells were transfected in 6-well tissue culture plates (Greiner) using Lipofectamine 2000 (Invitrogen) with 6 µg of target protein – GFP – 3xFLAG plasmid. After 24h the cells were lysed on ice using an Igepal lysis buffer (1% Igepal CA 630, 50 mM Tris-HCl pH 7.5, 150 mM NaCl, 10% Glycerol). Subsequently, the lysates were well mixed by pipetting the solution up and down for a minute, and the cell debris was pelleted by centrifuging the samples for 5 min. The supernatant was transferred and stored at −80°C.

### Design of AI-binders against immune-relevant proteins

An RFDiffusion pipeline similar to Watson et. al. ^6^ was used to generate binders to four intracellular targets. The crystal structure of PYCARD was obtained from the PDB (PDB: 7e5b) while the structures of p65, NLRP3, and RBM39 were obtained from the AlphaFold Protein Structure Database (https://alphafold.ebi.ac.uk/). Target structures were trimmed to just the domains of interest and hydrophobic “hotspot” residues were selected in the center of each interface. 10,000 diffusion trajectories were run against each target with several intermediate timesteps output in addition to the final timestep. ProteinMPNN ^7^ was used to assign sequences to each backbone (in triplicate) and Rosetta ddG ^24^ was used to filter the designs to the best 150,000 outputs per target. AlphaFold2 with initial guess ^25^ was used to predict the structures and pae_interaction ultimately used to rank them (with interface_rmsd < 4 and pae_interaction < 10 as absolute cuts). The 300 best interfaces for each target had their secondary structural elements extracted for additional resampling. RFDiffusion was then used to build 300 new backbones around each interface and those backbones were then subjected to 10 single-step partial diffusion trajectories to add diversity. ProteinMPNN and AF2 were used in a similar manner as the initial diffusion outputs to arrive at predicted binders. The top 500 binders to each target as ranked by AF2 pae_interaction and Rosetta ddG were selected for experimental testing.

### Re-scoring of AI-binder designs and mutants

Chai-1 (v.0.5.2), was used via the Python interface provided by the authors ^23^. The model was run with sequence information and ESM2 embeddings as input on a single Nvidia L40S GPU. No MSAs were provided to the model. For scoring, used the standard settings: 3 truck recycles, 200 diffusion time steps. Five predictions were used to calculate the median aggregate_score of each input sequence.

### AI-binder cloning

The binder oligo pools were designed not to contain relevant restriction sites and possess varying codons in the first bases of the binder, to reduce the necessary reads lengths for unambiguous determination. Furthermore, Esp3I restriction sites were added to each end of the binder. The binder oligo pool was amplified in a PCR reaction using binder binder_pool_fwd/rev primers and NEBNext PCR polymerase. After DNA purification, 100 ng backbone plasmid and 50 ng purified binder insert was added to a Golden Gate reaction that was running overnight. On the next day, the reactions were purified and concentrated using the Zymo DNA Clean & Concentrator-25 columns. The library was then electroporated, as previously described.

### Single AI binder cloning

Single AI binders were amplified using binder specific PCR primers and cloned into EGFP, mCherry, BTB and FKBP12 ^F36V^ backbones using Golden Gate reactions. Cloned constructs were transformed into Stable Competent *E. coli*. Transformants were sequence verified using Tn5-mediated whole-plasmid tagmentation and MiSeq sequencing.

### Generation of lentivirus of single AI-binders

1 × 10^6^ HEK293T wildtype cells were seeded in a 6-well one day prior to transfection to achieve ∼70% confluence the next day. For transfection, Lipofectamine2000 and OptiMEM serum-reduced medium were pre-incubated 5 minutes at RT. The lentiviral packaging plasmids 880 ng pMD2.G (Addgene #12259) and 1320 ng psPAX2 (Addgene #12260) and 1760 ng lentiviral binder vector were mixed in OptiMEM serum reduced medium. Lipofectamine and DNA mix were combined and incubated for 20 minutes at RT before dropwise addition to the HEK293T cells. After 4-6 hours the medium was replaced by DMEM GlutaMax medium supplemented with 10% FCS and 10 µg/mL Ciprofloxacin. 48 hours post-transfection the lentiviral supernatant was removed from the cells using a syringe and a blunt needle and filtered through a 0.45 µm filter. Filtered virus was either directly used for transduction of cells with 10 µg/mL polybrene, frozen in aliquots at – 80°C.

### Generation of reporter screening cell lines

HEK293T mCherry-2A-hASC-mCitrine cells were generated by retroviral transduction and subsequent fluorescence-based sorting. In brief, 1 × 10^6^ HEK293T cells were seeded per well one day prior to transfection. For transfection, a DNA mastermix was prepared in a final volume of 10 µl OptiMEM, containing 2 µg of the retroviral construct (pR mCherry-2A-hASC-mCitrine), 1 µg gag-pol, and 100 ng VSV-G. Separately, a transfection reagent mix was prepared by combining 8 µl GeneJuice with 100 µl OptiMEM and incubating at RT for 5 minutes. The DNA mastermix and transfection reagent mix were then combined and incubated for 20 minutes at RT before being added to the cells. Prior to transfection, the medium on the cells was replaced with 2 mL fresh DMEM complete medium. HEK293T cells were transfected with 110 µl of the combined DNA and transfection reagent mix. 16 hours post-transfection, the medium was exchanged for DMEM supplemented with 30% FBS. Thirty hours after this medium change, the retroviral supernatant was collected, filtered through a 0.45 µm syringe filter, and stored at −80°C. HEK293T cells were transduced with retrovirus in the presence of 10 µg/mL polybrene. Transduced cells were expanded until a T25 flask reached 90% confluency. Successfully transduced cells were sorted based on the desired fluorescence using a SONY MA-900 sorter with a 100 µm nozzle.

### Proteome manipulation assays in mammalian cells

For proteome manipulation assays using AI-binders in BTB-mCherry or dTAG-Puro vectors lentivirus of each AI-binder was generated as described above. HEK ASC-GFP or THP-1 wt cells were transduced with AI-binders in BTB-mCherry vectors and sorted for mCherry-GFP double-positive cells using a SONY MA-900 sorter with a 100 µm nozzle. To monitor induced protein aggregation 2 × 10^4^ cells per well were seeded in 96-well Carrier Ultra microscopy plate. Cells were stimulated with either 10 µM BI-3802 or DMSO for 16 hours. Protein aggregation was analyzed by confocal spinning disc microscopy. HEK mCherry-2A-hASC-mCitrine or THP-1 wildtype cells were transduced with AI-binders in dTAG-Puro vectors and puromycin selected for 3-5 days or until untransduced control cells were dead. For FACS-based detection of targeted degradation in HEK cells, cells were seeded in 96-well tissue culture plates with 2 × 10^4^ cells per well. The next day cells were stimulated with 10 µM dTAG-v1 or DMSO for 24 hours. Cells were detached and analyzed using a MACS Quant Flow Cytometer.

### Immunoblots of THP-1 cells

1 × 10^5^ cells per 96-well of THP-1 wildtype cells transduced with AI-binder dTAG-Puro constructs were differentiated at overnight in the presence of 100 ng/mL PMA. The next day cells were primed with 200 ng/mL LPS for 3 hours followed by 24 hours stimulation with 1 µM dTAG-v1. Control samples were stimulated with LPS only. After the stimulation, cells were gently washed with DPBS and lysed with 1x Lämmli buffer supplemented with beta-mercaptoethanol for 5 min at 95°C. Subsequently, proteins were separated on Mini Protean TGX Any kDa Stain Free gels and transferred to nitrocellulose membrane via wet blotting. Membranes were blocked using 2% milk in PBS-T for 1 hour, stained with 1 µg/mL murine anti-ASC (TMS-1) antibody (Biolegend) overnight at 4°C and secondary anti-mouse-HRP antibody (Invitrogen) for 1 hour at RT. Proteins were detected using Pierce ECL Western Blotting Substrate and imaged with a Fusion FX imager.

### NNK oligo pool design and library preparation

Oligo pools with one amino acid exchanges were designed by generating one DNA sequence per amino acid position that contains a codon consisting of mixed bases (NNK). Additionally, NNK oligos contain unique silent mutations around the randomized amino acid, to distinguish real mutations from sequencing errors. The oligo pools were generated with 30 bp Gibson cloning overhangs. A NEBNext Polymerase PCR with the oligo pool and the backbone vector was conducted. After amplification, Dpn1 was added to the backbone PCR for 30 minutes at 37°C to remove original plasmids from the mix. Both amplified fragments were added to a Gibson cloning reaction for 1 h at 50°C. The resulting library was purified and concentrated using the Zymo DNA Clean & Concentrator-25 columns and electroporated, as previously described. Transposon-Display was performed as previously described, with the exception that non-binder bacteria were included in the NNK pool at a final mixing ratio of 10:1.

### Neutralized binder generation

The NNK screen data was used to extract mutations that modify the binders to have a neutral charge at pH7.4 and include mutations that improve the binders’ performance in Transposon-Display. To validate the expression and binding capabilities of these modified neutral binders, smaller pools of binders with neutralizing amino acid mutation combinations were ordered as oligo pools, amplified with binder_pool_fwd/rev primers and cloned into the Transposon-Display vector using Golden Gate Assembly. Transposon-Display was performed as previously described, with the exception that non-binder bacteria were included in the neutral pool at a final mixing ratio of 10:1.

### Binder-fluorescent protein fusions

Single binder plasmids were retransformed, and a 2 ml culture in an Eppendorf tube with 1.5 ml LB-Amp medium was inoculated with one colony to grow overnight at 37°C. Bacterial cultures were used to inoculate 500 ml LB-AMP cultures until OD600 reached 0.4. Then, the bacterial cultures were induced with Anhydrotetracycline and on the next day sonicated, as described before. The bacterial lysate was aliquoted and stored at −20°C.

### AI-binder pull down

GFP-Catcher (Antibodies-Online) agarose beads were washed three times with PBS before adding them to the EGFP-binder bacterial lysate. After rotating for 2 hours at 4°C, the beads were pelleted in a precooled centrifuge and washed with ice cold PBS for 3 times. Afterwards, the beads were added to lysates of primary human macrophages or differentiated and stimulated THP1 cells. After rotating for 2 hours at 4°C, the samples were washed three times with pre-cooled 50 mM Tris-HCl pH 7.5 and after removing the buffer frozen at −80°C.

### Data-independent acquisition mass spectrometry proteomics

Pulldown samples were digested on beads using trypsin and peptide mixtures were analyzed using a Vanquish Neo liquid chromatography system (Thermo Fisher Scientific) coupled to an Orbitrap Astral mass spectrometer (Thermo Fisher Scientific). Raw files were analyzed using Spectronaut 19 (Biognosys).

### Cell staining

Primary human macrophages were generated and harvested from leukocyte-enriched buffy coats as described in Fandrey and Jentzsch et al. ^26^ Furthermore primary human macrophages were primed with 20 ng/mL LPS (InvivoGen) for 3h. To inhibit caspase-mediated cell-death pathways, the cells were incubated with 40 µM VX-765 (Invivogen) and 50 µM Z-VAD-FMK (MCE, #HY-16658B) for 30 minutes prior to stimulation. To initiate an inflammasome response, 1 µM PA (Biozol, LBL-171E) together with 10 ng LFn-PrgI were added to the cells. The cells were fixed in 4% PFA in PBS solution for 1 h and then permeabilized with 10% FCS and 0.1% TritonX-100 in PBS for 1 h. The EGFP-binder lysates were added to the fixed cells for 1 h. Afterwards, the cells were washed three times with PBS and stained with 2 µM Hoechst 33342. Cells were imaged with Z-stacks at a distance of 2.5 µm.

### Imaging

All images were acquired using a Nikon Ti2 body equipped with a Yokogawa CSU-W1 spinning disc unit connected to Lumencor Celesta multimode lasers with wavelengths of 405 nm (nuclear staining), 477 nm (GFP), and 546 nm (mCherry). Emission filters used were Chroma ET450/50 (nuclear staining), Chroma ET525/50 (GFP), 572/28 BrightLine HC (mCherry). The exposure time was 90 ms for all channels. Objectives used were Nikon 20x CFI P-Apo with a 1.5x tube lens inserted into the light path. A Hamamatsu Orca Flash 4.0 LT+ camera was used in electronic shutter mode at full resolution (2048×2048).

## Data availability

NGS data and microscopy datasets will be available upon request.

## Code availability

Custom design and analysis tools will be available as open-source web applications at *www.jsb-lab.bio/AI-binders*.

